# An empirical Bayes method for serotype case-carrier ratios, with an application to Group B streptococcus

**DOI:** 10.1101/421412

**Authors:** Joseph A. Lewnard, Lauren A. Cowley

**Author notes:** Corresponding author: Joseph Lewnard, PhD, Division of Epidemiology & Biostatistics, School of Public Health, University of California, Berkeley, 2121 Berkeley Way #5412, Berkeley, California 94720. **Conflicts of interest:** JAL declares receipt of research grants from Pfizer to the University of California, Berkeley and to Harvard University for unrelated work.

## Abstract

**Background:** Case-carrier ratios quantifying the relative pathogenicity of serotypes can inform vaccine formulations for antigenically-diverse pathogens. However, sparse serotype-specific counts in epidemiologic datasets may undermine such analyses, most notably for rare serotypes that pose emergence risks in vaccinated populations. This challenge is well-illustrated in Group B streptococcus (GBS), where serotype III dominates in both carriage and disease.

**Methods:** We develop an empirical Bayes random-effects model based on conjugate Dirichlet-multinomial distributions of serotype frequencies in carriage and disease states. We validate the model using simulated datasets, and apply it to data from 15 paired sets of GBS isolates from intrapartum rectovaginal colonization (n=3403) and neonatal invasive disease (NID; n=1088), 16 from blood (n=2352) and cerebrospinal fluid (n=780) neonatal specimens, and 3 from fatal (n=173) and non-fatal (n=1684) neonatal invasive infections.

**Results:** Our method accurately recovers parameters in simulated datasets. Using this approach, we confirm that GBS serotype III exhibits the greatest invasiveness, followed by serotype Ia with a 75.3%
(95%CrI: 43.7-93.8%) lower estimate. Enhanced invasiveness of serotypes III and Ia is most evident in late-onset disease. Non–hexavalent-vaccine serotypes, which are rare in carriage and disease, generally show lower invasiveness; serotype IX/non-typeable GBS, the most prevalent cause of non–vaccine-preventable disease, is 98.7% (81.7-99.9%) and 94.2% (13.9-99.6%) less invasive than serotypes III and Ia, respectively.

**Conclusions:** We present a strategy for measuring associations of serotype with carrier and disease states in the presence of sparse counts, avoiding biases that exist in common ad-hoc approaches.

## INTRODUCTION

Group B streptococcus (GBS; *Streptococcus agalactiae*) is the leading cause of neonatal invasive infections, with 0.53 cases per 1000 live-births globally^1^ and 2.03 cases per 1000 live-births in subsaharan Africa.^2^ While intrapartum antimicrobial prophylaxis prevents early-onset GBS disease (EOD; occurring 0-6 days after birth),^3^ additional preventive strategies are needed to combat late-onset disease (LOD; 7-90 days), and to serve populations with limited access to antenatal care. There are ten recognized serotypes of GBS defined by distinct polysaccharide capsular antigens (Ia, Ib, II-IX) as well as non-typeable strains (NT-GBS). Anticapsular conjugate vaccines intended for pregnant women are currently under development. A trivalent conjugate vaccine (GlaxoSmithKline) targeting serotypes Ia, Ib, and III has completed phase-2 trials,^4,5^ and a hexavalent vaccine (Pfizer) against serotypes Ia, Ib, and II-V is entering human studies.

Considerations around GBS vaccine compositions have centered around serotype frequencies in neonatal invasive disease, with serotypes Ia, Ib, and III alone comprising 85% of cases worldwide.^6^ However, concerns surround the possibility that vaccine-derived immunity against targeted GBS serotypes may exert selective pressure on the circulating GBS population, enabling non-vaccine serotypes that are currently rare to expand in prevalence.^7^ Such “serotype replacement” in *Streptococcus pneumoniae*—whereby serotypes not targeted by pneumococcal conjugate vaccines (PCVs) have overtaken the ecological niche formerly inhabited by vaccine-targeted serotypes—has offset the impact of PCVs against invasive pneumococcal disease.^8^ Monitoring for serotype replacement has also been central to postlicensure surveillance of the impact of vaccines against human papillomavirus,^9^ meningococcus,^10^ and *Haemophilus influenzae* type b.^11^

Estimates of the relative capacity of serotypes to cause infections, severe clinical manifestations, or death provide a basis for forecasting the impact of vaccines on the epidemiology of antigenically-diverse pathogens. Experience with PCVs has illustrated that these estimates can lead to better vaccine compositions than historical approaches,^12^ such as the selection of the most prevalent serotypes as vaccine targets,^13,14^ under the scenario of serotype replacement. Such estimates also provide a baseline for monitoring post-vaccination epidemiological changes.^15,16^ While historical data suggest enhanced invasiveness of GBS serotype III relative to serotypes Ia, Ib, and II, these studies pre-date the emergence or discovery of serotypes IV-IX and nontypeable GBS (NT-GBS) lineages, and are of limited size, statistical rigor, and geographic scope.^17–19^ Thus, quantifications of the relative pathogenicity of GBS serotypes are necessary to assess the potential impact of anticapsular vaccine candidates.

The rare occurrence of certain GBS serotypes in maternal carriage and neonatal disease, however, makes it difficult to estimate these quantities with certainty, even in large samples.^20,21^ While random-effects meta-analysis of data from multiple epidemiologic studies may thus present an attractive analysis strategy,^22^ investigators remain likely to encounter challenges when generating pooled risk ratios or odds ratios in the presence of zero-valued counts within studies. Common ad-hoc workarounds, such as “continuity corrections” adding a fixed value to cells^23–25^ and simple summing or averaging of values across studies,^13^ risk introducing bias in both point estimates and measures of uncertainty.^26^ While many studies exclude analyses of rare serotypes to avoid such problems,^27–31^ rare serotypes may in fact be of interest for serotype replacement; moreover, zero-valued cell counts within studies present valuable information on overall serotype prevalence. Here, we develop an empirical Bayes method to infer serotype distribution parameters in pooled analyses of sparse datasets, providing a basis for estimating case-carrier ratios. We apply this approach to estimate the association of GBS serotype with invasive potential, risk of presenting as meningitis, and risk of death in neonatal infections.

## METHODS

### Outcome definition

We aimed to measure the association of serotype with the following epidemiologic features of neonatal invasive GBS:

1. Risk of causing invasive disease given maternal colonization: We defined invasive disease as GBS isolation from an ordinarily sterile site (e.g., blood or cerebrospinal fluid), distinguishing between EOD and LOD;
2. Risk for meningitis during neonatal invasive infection: We defined cases with GBS isolation from cerebrospinal fluid as meningitis; and
3. Risk for fatal outcome of neonatal invasive infection: We defined fatal cases as deaths attributable to invasive GBS among live-born infants (thus excluding stillbirths).

For analyses of invasive potential, we compared serotype frequencies in neonatal disease against serotype frequencies in recto-vaginal colonization among pregnant women; we did not restrict colonization samples according to time of sampling within pregnancy due to limited metadata on when swabs were obtained.

### Study identification

We updated previously-undertaken systematic literature reviews^1,6^, and extracted serotype-stratified data on maternal carriage, disease timing (EOD/LOD), disease manifestation (meningitis or other), and outcome (fatal or non-fatal). For analyses of serotype associations with neonatal invasive potential, we aimed to include all studies monitoring both carriage and disease serotype distributions within a single source population or setting; for analyses of serotype associations with meningeal invasion and fatal outcome, we sought all studies presenting serotype frequencies in neonatal meningitis and sepsis, and in fatal and non-fatal neonatal invasive disease. We examined full texts of all studies included in the previous reviews to determine eligibility, and conducted a PubMed search as well as citation tracking of these reviews and two other relevant studies^20,21^ to identify new articles for screening. We detail the search strategy and inclusion/exclusion criteria in the **Supplementary information**.

### Statistical framework: overview

From each study with paired carriage and disease data, we estimated the relative odds of a serotype being isolated from neonatal invasive disease (all neonatal invasive disease, EOD, LOD) versus maternal colonization; we derive this measure on the risk scale below. From studies reporting GBS isolation from blood or CSF, we estimated the relative risk that invasive disease caused by each serotype would manifest as meningitis. Last, for studies reporting fatal and non-fatal outcome, we estimated the risk ratio for fatal outcome (case-fatality ratio; CFR), given invasive disease, for each serotype.

We pooled these measures of association across studies under an empirical Bayes framework, described below in technical detail. Briefly, we took serotype frequencies in carriage, all neonatal invasive disease, EOD, LOD, meningitis, sepsis, non-fatal disease, and fatal disease, within each study, to be multinomially-distributed. We took the parameters of each setting-specific multinomial distribution, in turn, to be random draws from a global Dirichlet conjugate prior distribution. This enabled us to use data from all studies in estimating Dirichlet hyperparameters describing global serotype frequencies in GBS carriage and disease, and thus to generate positive-valued estimates of setting-specific multinomial probabilities in the presence of zero-valued counts. In turn, we used these multinomial probabilities to generate pooled measures of association under a random effects model.

### Empirical Bayes random effects model

The invasive potential of serotype *j*, measured relative to serotype *k*, from serotype frequencies in carriage and disease (**x**^*C*^ and **x**^*D*^, respectively), is typically estimated as

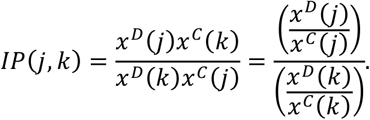

We derive this in a later section (“**Measures of association**”). Challenges arise when **x**^*C*^ and **x**^*D*^ include zero-valued counts. We propose the following solution: Within the *ℓ*^th^ study, take observed serotype-specific frequencies in carriage and disease (vectors 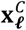 and 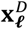, respectively) to be random draws from a multinomial distribution, specified in full by the number of carriage and disease isolates obtained (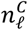 and 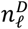, respectively) and the true underlying proportions of carriage and disease (vectors 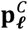 and 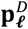, respectively) attributable to each serotype within the populations from which cases and carriers were sampled:

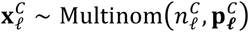

and

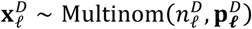

The Dirichlet conjugate prior for the population proportion parameters, 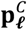 and 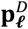, defines these study-level proportions as random deviates of global serotype distributions with parameters **α**^*C*^ and **α**^*D*^:

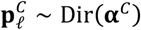

and

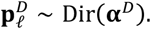

This yields positive estimates of 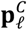 and 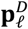 across the included studies. We verified the identifiability of the parameters in a simulation study described below.

Define study-level estimates of neonatal invasive potential using the inferred population proportions of carriage and disease attributable to each serotype. Within the *ℓ*^th^ study,

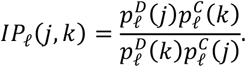

We calculate invasive potential relative to serotype III (*k*), taking measures from each study to be deviates from a global serotype-specific value, *q*(*j,k*), with associated variance across studies *σ*^2^(*j,k*):

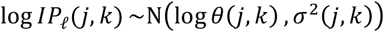

### Estimation

We sampled from the joint distributions 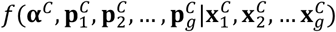 and 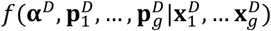 via Markov chain Monte Carlo (MCMC) using a Metropolis-Hastings updating procedure. We ran chains for 600,000 iterations after the burn-in period, saving the state of the chain at every 500^th^ sample to reduce autocorrelation.

We estimated the global invasiveness parameters according to

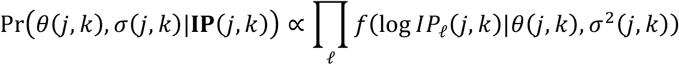

which provided a basis for sampling from the joint distribution of {*θ*(*j, k*), *σ*^2^(*j, k*)} for each serotype *j*, relative to serotype III (*k*).

### Measures of association

#### Odds ratio estimator for neonatal invasive potential

Define the neonatal invasive potential (IP) of serotype *j* relative to serotype *k* as the ratio

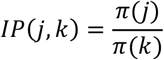

of the probabilities *π*(*j*) and *π*(*k*) for neonates exposed to maternal carriage of serotypes *j* and *k*, respectively, to experience invasive disease, such that

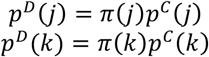

for the variables **p**^*C*^ and **p**^*D*^ defined above. We note that *π* represents the joint probability of both GBS transmission from mother (or other source) to neonate and of disease progression given transmission. Thus,

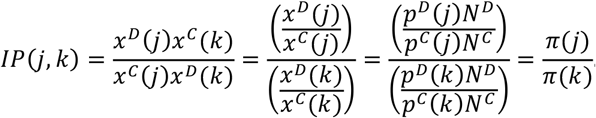

#### Risk ratios for meningitis and fatal outcome

The conventional relative risk estimator for invasive disease cause by serotype *j* to manifest as meningitis, as compared to invasive disease caused by serotype *k*, is

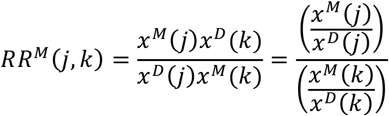

for the vectors of serotype frequencies in meningitis (**x**^M^), here specifying that **x**^*D*^ includes cases with known clinical manifestations. Similarly, the relative risk for invasive disease caused by serotype *j* to result in fatality, as compared to invasive disease caused by serotype *k*, is conventionally measured as

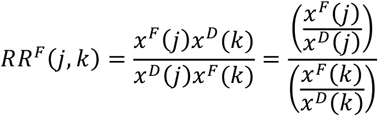

where **x***^F^* indicates serotype frequencies in fatal cases of illness, while **x***^D^* includes cases with known clinical outcomes (fatal or nonfatal). Consistent with the approach taken for invasive potential, we sampled from the distributions of serotype-specific proportions in all meningitis cases (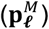) and fatal invasive disease cases (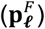) via

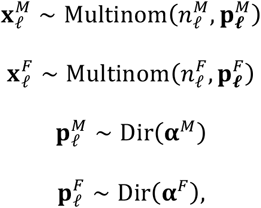

where 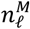 and 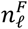 indicate the total number of meningitis isolates and fatal case isolates, respectively, in the *ℓ*^th^ study, and **α***^M^* and **α***^F^* serve as hyperparameters for the underlying distributions. We define global estimates for serotype-specific relative risks from the normal distribution of log-transformed estimates from each study as above:

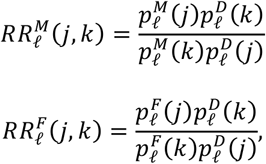

and use the same framework to estimate *θ*(*j, k*) and *σ*(*j, k*) as we describe above for the analysis of invasive potential.

#### Presentation

For all analyses, we report relative measures (odds ratios and risk ratios) using the serotype with the greatest absolute odds or risk of the endpoint as the referent category.

### Simulation study

We conducted a simulation study to verify identifiability of the **p***_ℓ_* parameters, defined above, that we used as inputs to calculation of case-carrier ratios. We generated simulated distributions of 10 serotypes from the Dirichlet conjugate prior distribution. We considered various parameterizations to allow scenarios of differing serotype diversity, defining *α*(*j*) as exp (−*λk*(*j*))*b*, with **k** comprising randomly-ordered values of 1,2, …, 10. We considered *λ* ∈ (0.1,0.2, …,0.6) to allow differing degrees of serotype dominance, and *b* ∈ (1,5,10) to allow differing degrees of between-study variance in serotype-specific prevalence among isolates. From each of the resulting Dirichlet parameterizations for **α**, we sampled twenty vectors **p***_ℓ_* for differing study settings, and generated observations

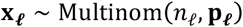

with sample sizes *n_ℓ_* distributed uniformly, on the log scale, between 10 and 1000. We used the simulated data **x**_*ℓ*_ to estimate **p***_ℓ_* and **α** via Markov chain Monte Carlo sampling, consistent with the approach described above. We calculated parameter coverage probabilities at the 2.5% credible threshold (an analog to statistical power for continuous measurements, indicating the probability that true values of these parameters were contained within 95% credible intervals of the estimates).

## RESULTS

### Model performance on simulated data

We illustrate the proportion of counts expected to take on values of zero under differing values of *b* and *l* in **Figure 1A-C**, for sample sizes between 10 and 1000 isolates. Zero-valued counts are most probable in small samples, at low values of *b* (corresponding to greater variance across settings in multinomial serotype prevalence), and at high values of *λ* (corresponding to a situation where one or several serotypes is consistently dominant among carriage or disease isolates).

**Figure 1:**
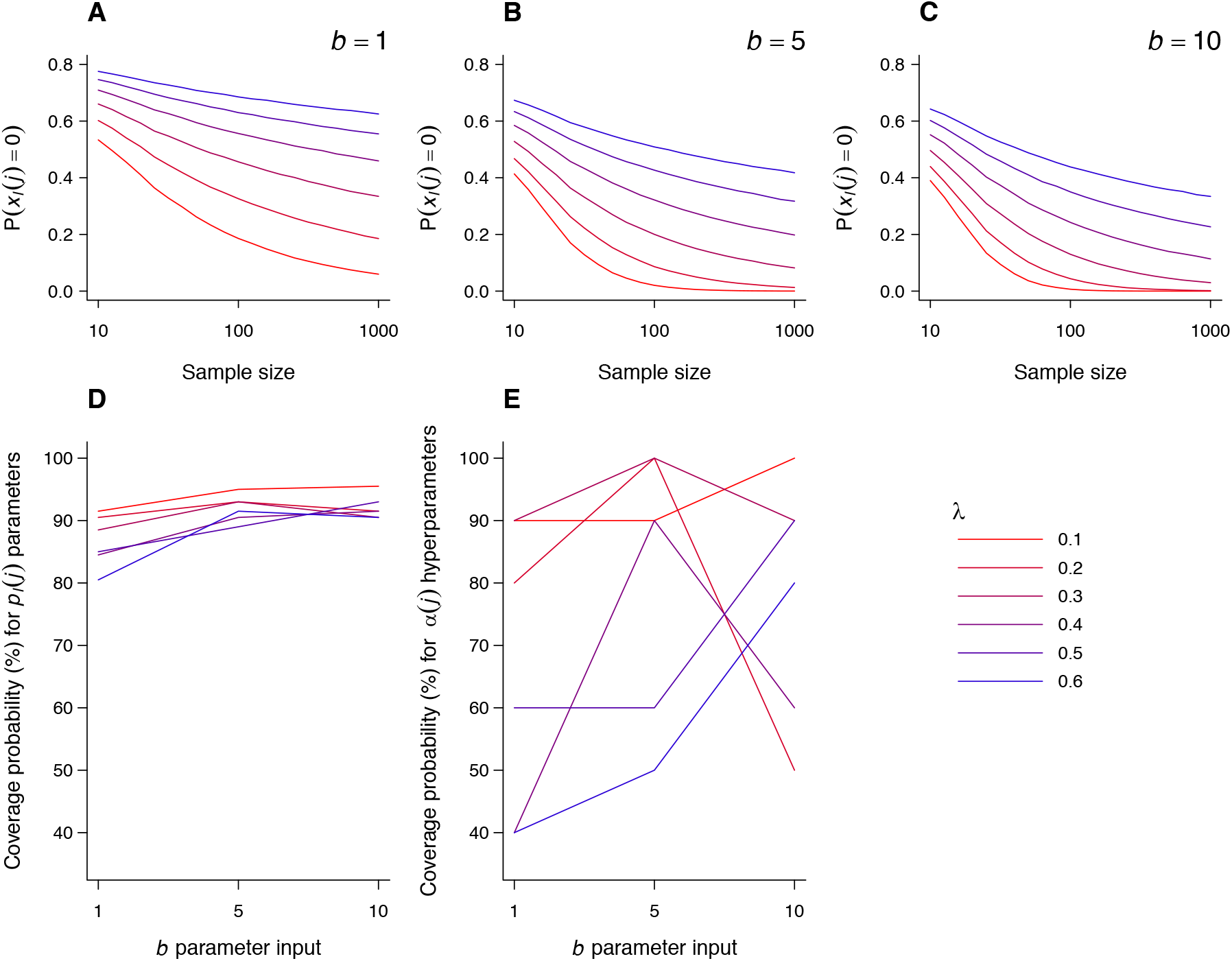
Assessment of parameter identifiability in simulated data. A-C) To illustrate the range of epidemiologic situations covered by our simulation study, we illustrate the probability for serotype-specific counts in individual studies (*x_ℓ_*(*j*) for count *x* of serotype *j* in study *ℓ*) to take on values of zero under differing sample sizes (x-axis) and values of the parameters *b* and *λ*. Higher values of *b* correspond to lower variance across settings in the prevalence of a given serotype prevalence among GBS isolates *λ*. Higher values of *b* correspond to a scenario where a given serotype (or serotypes) occupies a greater proportion of all isolates, i.e. reduced Simpson diversity among serotypes. D) We identify 80% or greater coverage (probability for 95% credible intervals to contain the true value) for estimates of **p**_*ℓ*_, the setting-level prevalence of serotypes among all isolates, under the conditions explored. Estimates approach 95% at increasing values of *b* and decreasing values of *λ*. E) We identify poorer coverage for estimates of the hyperparameter **α**, which also tend improve under the conditions that lead to more accurate estimation of **p**_ℓ_; our method does not apply the hyperparameter toward estimates of epidemiologic association.

We identified 80% or greater probability for true estimates of setting-specific serotype prevalences (**p**_*ℓ*_) to fall within 95% credible intervals of estimates across the range of values explored for *b* and *λ* (**Figure 1D**); these coverage probabilities were poorest under the circumstances that would lead to the greatest probability for zero-valued counts as described above (low *b*, high *λ*). Simulated coverage probabilities for **p**_*ℓ*_ fell within the range of 80.5-91.5% for *b*=1, and 90.5-95.5% for *b*=10. A similar trend toward better coverage probabilities with higher *b* and lower *l* was evident in estimates of the hyperparameter **α** (**Figure 1E**); lower accuracy in estimates of **α** (with coverage probabilities reaching as low as 40%) did not prevent the model from recovering valid estimates of the target parameters **p**_*ℓ*_.

### Data abstraction

We assessed full texts of 599 studies covered in previous systematic reviews, and screened abstracts of 50 articles identified via a PubMed search and 529 articles citing the previous reviews and two other relevant articles, not exclusive of duplicates (**Figure 2**). From these, we identified 15 paired datasets of GBS serotype distributions in carriage and neonatal invasive disease (3403 and 1088 isolates, respectively; **Table 1**); 16 datasets presenting serotype distributions in meningitis and other neonatal invasive disease (3132 isolates; **Table 2**); and 3 datasets presenting serotypes isolated from fatal and non-fatal neonatal invasive disease cases (1684 isolates; **Table 2**).

**Figure 2:**
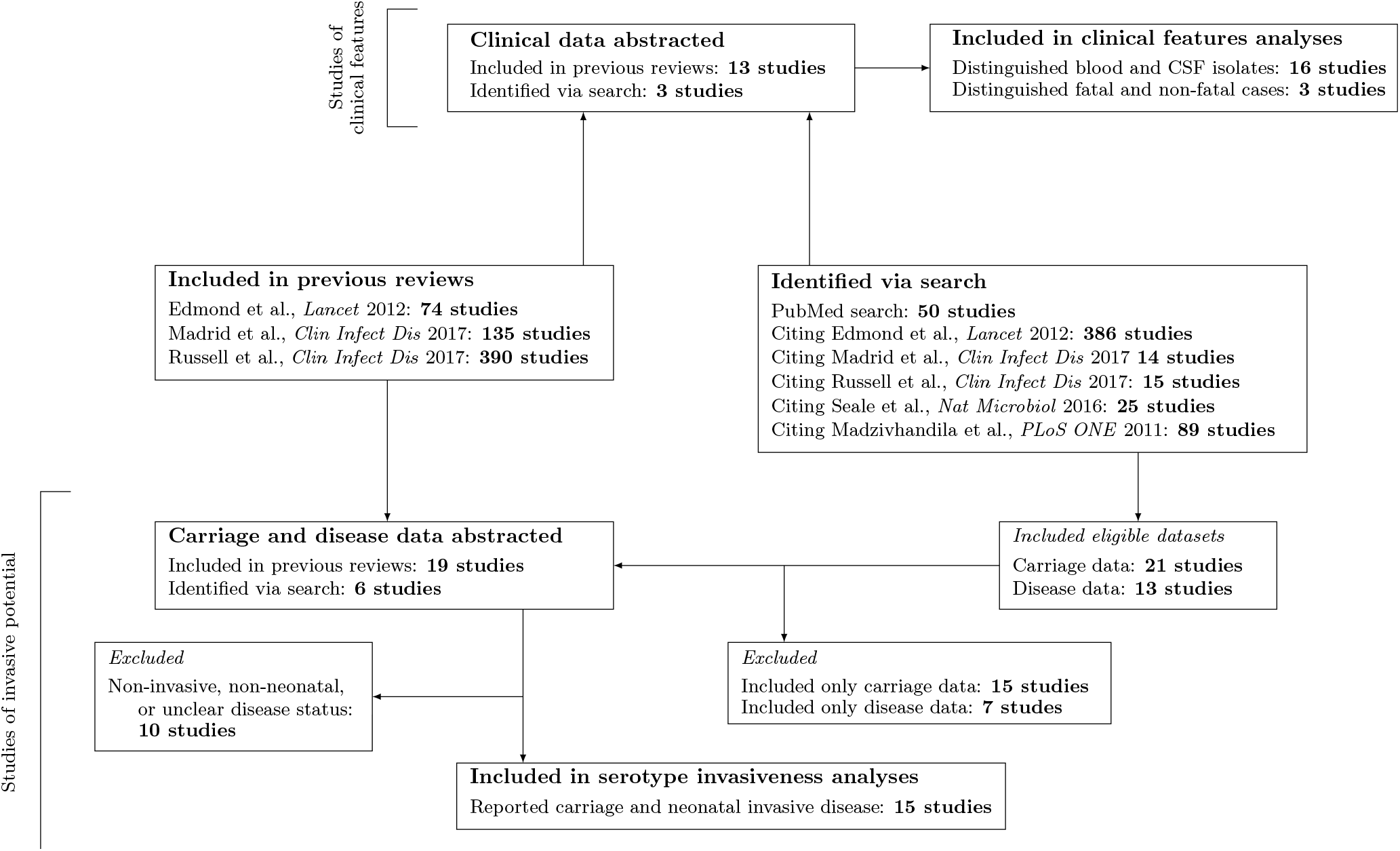
Systematic review flowchart. We detail the identification and inclusion/exclusion of articles from previous systematic reviews and articles as well as their citations, and a PubMed search for articles published after January 1, 2016. Tabulations do not exclude duplicate results.

**Table 1:** Included studies indicating serotype association with invasive potential.

| Study | Reference<br>(Supplement) | Setting | Basis for pairing | Isolates obtained |  |  |  |
| --- | --- | --- | --- | --- | --- | --- | --- |
|  |  |  |  | Carriage | Neonatal<br>disease | EOD | LOD |
| Ko et al., 2015 | 47 | Australia | Geography, date | 408 | 108 | 60 | 48 |
| Seale et al., 2016 | 48 | Kenya | Mother-infant<br>dyads | 915 | 70 | 30 | 40 |
| van Elzakker et al., 2009 | 49 | Taiwan | Geography, date | 58 | 43 | — — | — — |
| van Elzakker et al., 2009 | 49 | Netherlands | Geography, date | 92 | 44 | — — | — — |
| Madzivhandila et al.,<br>2011 | 50 | South<br>Africa | Mother-infant<br>dyads | 541 | 245 | 137 | 108 |
| Martins et al., 2007 | 51 | Portugal | Geography, date | 269 | 64 | 42 | 22 |
| Bisharat et al., 2005 | 52 | Israel | Geography, date | 104 | 50 | — — | — — |
| Berg et al., 2000 | 53 | Sweden | Geography, date | 114 | 128 | 40 | 10 |
| Moraleda et al., 2018 | 54 | Morocco | Geography, date | 67 | 5 | — — | — — |
| Lopez et al., 2018 | 55 | Spain | Facility, date | 242 | 79 | — — | — — |
| Figueira-Coelho et al.,<br>2004 | 56 | Portugal | Facility, date | 108 | 17 | — — | — — |
| Davies et al., 2001 | 57 | Canada | Geography, date | 118 | — — | 75 | — — |
| Teatero et al., 2014 | 58 | Canada | Facility, date | 102 | 93 | 34 | 59 |
| Martinez et al., 2004 | 59 | Chile | Geography, date | 65 | 10 | — — | — — |
| Hayes et al., 2017 | 60 | Ireland | Geography, date | 200 | 7 | — — | — — |

**Table 2:** Included studies indicating serotype association with disease manifestation and fatal outcome.

| Study | Reference<br>(Supplement) | Setting | Isolates obtained |  |  |
| --- | --- | --- | --- | --- | --- |
|  |  |  | Meningitis | Non-<br>meningitis | Fatal<br>outcome |
| Davies et al., 2001 | 57 | Canada | 17 | 74 | 91 |
| Ekelund et al., 2004 | 61 | Denmark | 107 | 405 | -- |
| Gray et al., 2007 | 62 | Malawi | 21 | 25 | -- |
| Zhao et al., 2008 | 63 | Australia, New<br>Zealand | 20 | 73 | -- |
| Martins et al., 2007 | 51 | Portugal | 4 | 55 | -- |
| Trijbels-Smeulders et al.,<br>2006 | 64 | Netherlands | 63 | 135 | -- |
| Fluegge et al., 2005 | 65 | Germany | 67 | 225 | -- |
| Chang et al., 2014 | 66 | Japan | 37 | 21 | -- |
| Liu et al., 2015 | 67 | Chiu | 3 | 5 | -- |
| Morozumi et al., 2014 | 68 | Japan | 81 | 48 | -- |
| Six et al., 2016 | 69 | France | 4 | 12 | -- |
| Teatero et al., 2014 | 58 | Canada | 9 | 80 | -- |
| Yoon et al., 2015 | 70 | Republic of Korea | 25 | 24 | -- |
| Siguaque et al., 2018 | 71 | Mozambique | 14 | 21 | -- |
| Dangor et al., 2016 | 72 | South Africa | 172 | 424 | 596 |
| Phares et al., 2008 | 73 | United States | 156 | 841 | 997 |

Of the paired maternal carriage and neonatal disease datasets, two were obtained from maternal-newborn dyads; all others were from geographically- and temporally-matched samples of maternal colonization and neonatal invasive disease isolates. Latin American (1 study, 75 isolates), East Asian (1 study, 101 isolates), and South/Southeast Asian (0 studies) populations were under-represented; nine datasets were from North America, Europe, and Oceania. Studies reporting disease manifestation and fatal outcome were from North America, Europe, Oceania, East Asia, and sub-Saharan Africa; no data from Latin America or South/Southeast Asia were available.

### Relative neonatal invasive potential across GBS serotypes

Serotype III caused 48-95% of neonatal invasive disease cases across the included studies, followed by serotype Ia (1-28%), serotype V (0-15%), and serotype II (0-10%). This hierarchy of serotype frequencies was less evident in colonization (**Figure 3**). Serotypes Ia, Ib, and II-VII and IX/NT-GBS were each associated with lower neonatal invasive potential in comparison to serotype III; because no studies reported serotype VIII in disease, we did not generate pooled estimates for this serotype. Relative to serotype III, we estimated serotypes Ia and V were 24.7% (6.2-56.3%) and 5.6% (0.9%-21.8%) as likely, respectively, to cause neonatal invasive disease given maternal carriage. These pooled estimates were consistent with findings of individual studies; nine and eleven studies, respectively, provided evidence of lower neonatal invasive potential of serotypes Ia and V (relative to serotype III) at the 97.5% confidence threshold. We estimated the neonatal invasive potential for serotypes Ib, II, and IV—the remaining serotypes included in the candidate hexavalent vaccine—to be 3.9% (0.4-17.4%), 2.9% (0.4-11.6%), and 8.6% (1.0-70.5%) that of serotype III, respectively, and 12.2% (1.6-70.5%), 16.1% (1.5-97.5%), and 36.8% (3.7–459.5%) that of serotype Ia.

**Figure 3:**
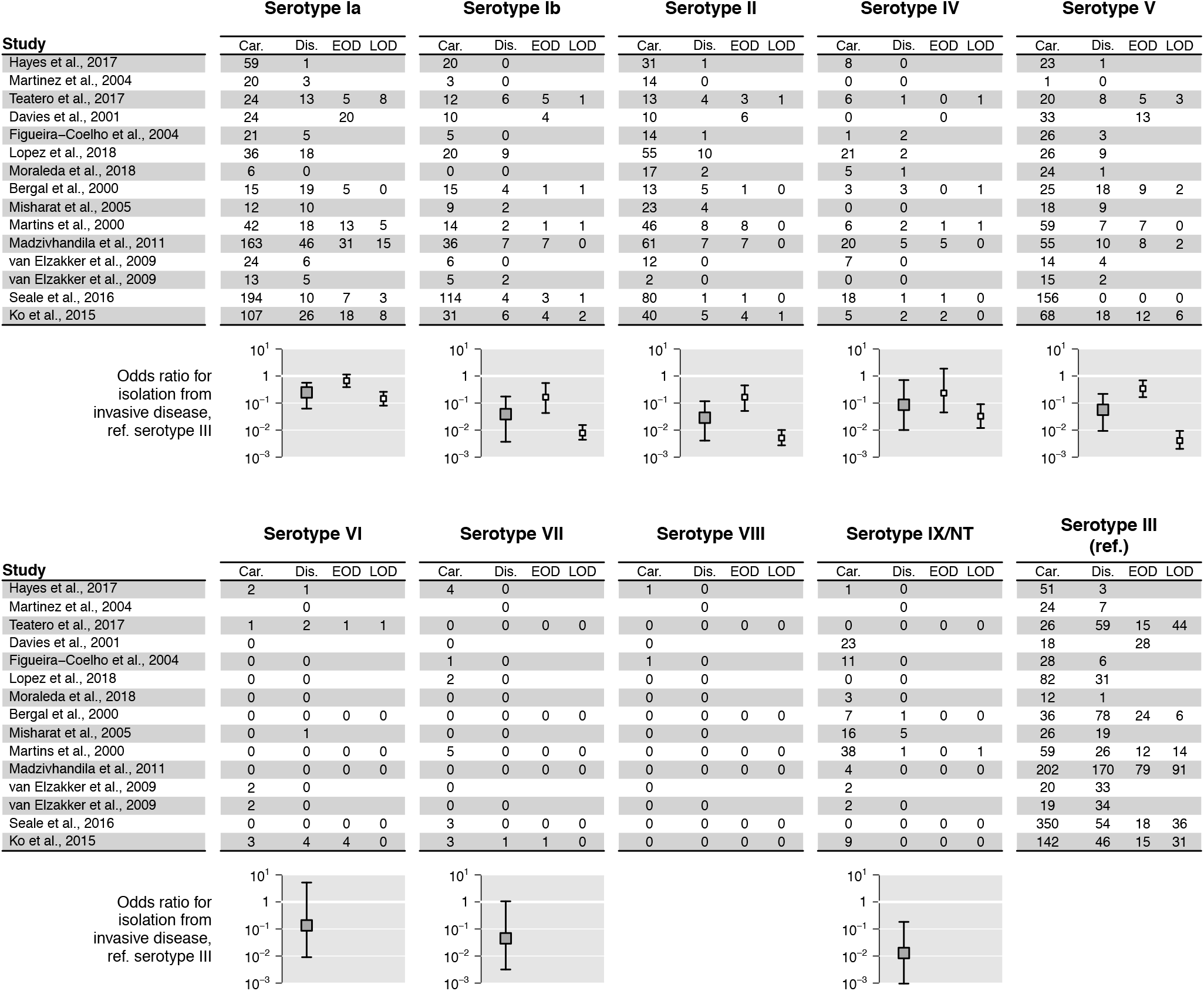
Association of serotype with neonatal invasive potential. For each study, we indicate serotype frequencies in intrapartum recto-vaginal colonization (“Car.”) and neonatal invasive disease (“Dis.”), as well as EOD and LOD, as permitted by reporting within original studies. We plot pooled estimates of serotype-specific relative invasive potential beneath serotype frequencies for all disease (grey points) as well as EOD and LOD (white points), measured from the odds ratio that an isolate came from invasive disease versus colonization, relative to serotype III (lower right corner). Our derivation of this measure is provided in the **Supporting information**. Pooled estimates include data from studies reporting only partial selections of serotypes, where total numbers of included isolates provide a basis for sampling from proportions attributable to each serotype. Bars signify 95% credible intervals around pooled estimates.

Serotype VI caused only 0-2% of neonatal invasive disease within studies. We identified 87.9% posterior probability that serotype VI was associated with reduced serotype potential relative to serotype III, but found its invasiveness to be statistically indistinguishable from that of serotype Ia at levels of statistical power afforded by our meta-analysis; we identified ≥;80% probability that serotype VI invasiveness exceeded that of vaccine-targeted serotypes Ib and II. We estimated the neonatal invasive potential of serotype VII to be 4.5% (0.3-105.1%) that of serotype III. Serotype IX/NT-GBS was associated with the lowest neonatal invasive potential, at 1.3% (<0.1-18.3%) that of serotype III and 5.8% (0.4-86.1%) that of serotype Ia; moreover, we identified ≥80% posterior probability that this reflected lower neonatal invasive potential than each serotype except Ib, II, and VII.

Serotype differences in invasive potential were less apparent in EOD than in LOD. Point estimates of EOD potential again suggested exceptional invasiveness for serotype III; we estimated the EOD potential of serotypes Ia and IV to be 67.0% (38.7-113.3%) and 23.5% (4.5-186.8%) that of serotype III, respectively, whereas these serotypes showed 14.6% (9.0-25.9%) and 3.3% (1.2-9.1%) the LOD potential of serotype III. For serotypes Ib, II, and V, we estimated EOD potentials to be 16.5% (4.3-54.9%), 16.5% (5.1-44.9%), and 34.1% (16.5-68.7%) as high as serotype III, respectively, whereas the LOD potentials of these serotypes were 0.8% (0.4-1.5%), 0.5% (0.3-1.0%), and 0.4% (0.2-0.9%) that of serotype III. Low serotype-specific frequencies in both carriage and disease (beneath the threshold at which our simulation study predicted reliable coverage) prohibited us from distinguishing EOD and LOD potentials for the non-vaccine serotypes.

### Relative risks of meningitis and death across GBS serotypes

Meningitis occurred in 10–77% of serotype III neonatal infections across the included studies; as compared to serotype III, serotypes Ia, Ib, II, and V showed 10-13% as high a risk of meningitis, given invasive disease (**Figure 4**). Serotypes IV, VI, and IX/NT-GBS were isolated in a lower proportion of studies, and caused meningitis in 0-100% of invasive disease episodes within these studies. Pooled estimates suggested they were each less likely than serotype III to manifest as meningitis in invasive disease cases, albeit with weaker precision as compared to estimates for serotypes Ia, Ib, II and V. Although we do not stratify estimates by EOD/LOD, we present data in **Figure 4** from studies that defined serotype frequencies within these strata.

**Figure 4:**
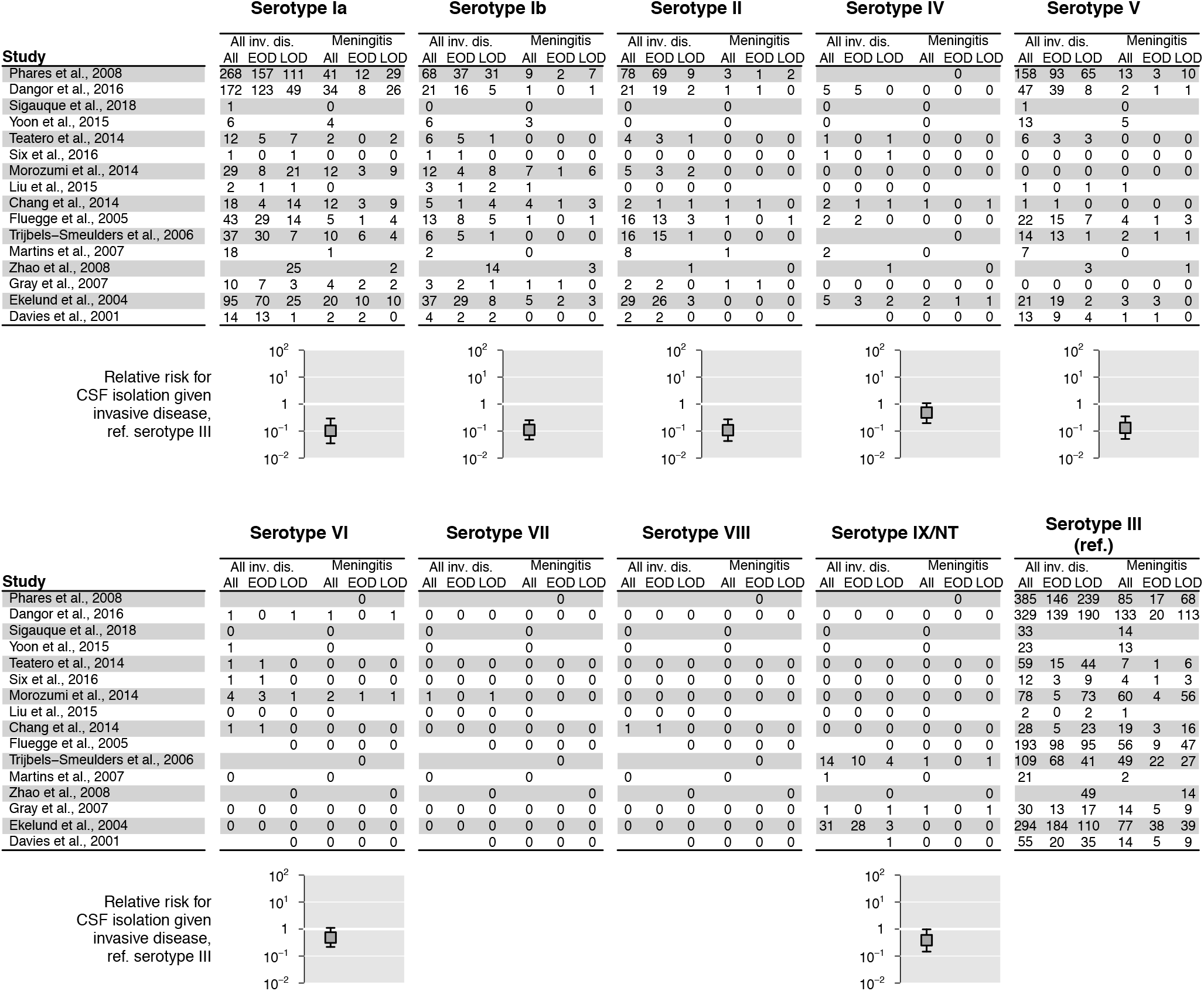
Association of serotype with meningeal invasion. For each study, we indicate serotype frequencies in all neonatal disease and in meningitis (signified by isolation from CSF), including all neonatal disease as well as EOD and LOD, as permitted by reporting within original studies. We plot pooled estimates of serotype-specific relative risk of meningitis beneath serotype frequencies for all disease (grey points), measured from the relative risk that an isolate came from CSF, measured against serotype III (lower right corner), for all infections. Our derivation of this measure is provided in the **Supporting information**. Pooled estimates include data from studies reporting only partial selections of serotypes, where total numbers of included isolates provide a basis for sampling from proportions attributable to each serotype. Bars signify 95% credible intervals around pooled estimates.

Risk of death showed lower degrees of variation among serotypes, with no estimated differences meeting conventional thresholds of statistical significance (**Figure 5**). Point estimates suggested the highest CFR for neonatal invasive disease caused by serotype Ia, similar to the estimtes for serotype V and VI with relative risks of 90.5% (15.7-376.2%) and 77.9% (5.0-902.5%), respectively. We estimated the CFR for disease caused by serotype III to be 46.1% (5.6-271.8%) that of serotype Ia; point estimates for the relative risk of death given invasive infection were in the range of 31-59% for the remaining vaccine-targeted serotypes. No fatal cases of disease caused by serotypes VII, VIII, and IX/NT-GBS were reported in the original studies.

**Figure 5:**
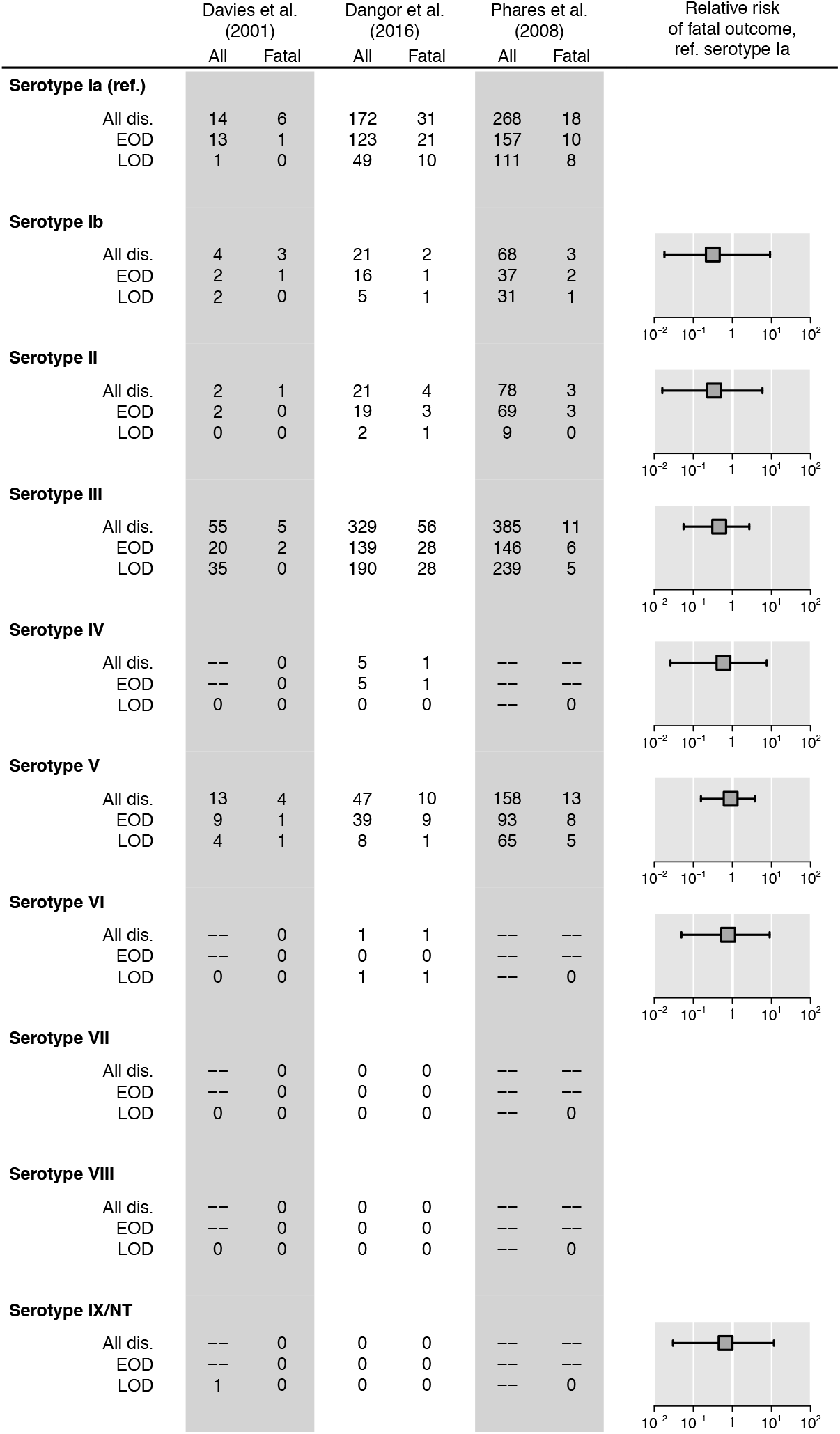
Association of serotype with risk of fatal outcome. For each study, we indicate serotype frequencies in all neonatal invasive disease and in fatal cases, including totals as well as EOD and LOD. We plot pooled estimates of serotype-specific relative risk of death given invasive infection, measured against serotype Ia, for all infections. Pooled estimates include data from studies reporting only partial selections of serotypes, where total numbers of included isolates provide a basis for sampling from proportions attributable to each serotype. Bars signify 95% credible intervals around pooled estimates.

## DISCUSSION

Because vaccines targeting antigenically-diverse pathogens may facilitate expansion of non-vaccine serotypes, assessments of serotype-specific pathogenicity provide key insight into the potential impacts of serotype replacement on disease burden. We present a method that circumvents key limitations to such analyses. These include the need to limit such analyses to only the most prevalent serotypes due to zero-valued counts of serotype frequencies in datasets, and to implement ad-hoc continuity corrections that bias measures of statistical association and uncertainty.^26^ Because non-vaccine serotypes posing emergence risks in vaccinated populations may be uncommon in the pre-vaccine era, unbiased quantification of rare serotype parameters is of particular importance.^12^ Our empirical Bayes approach uses estimates of serotype-specific prevalence across included studies to generate a “global” Dirichlet conjugate prior for serotype prevalence among GBS isolates in study populations, treating the observed serotype frequencies within each setting, in turn, as independent multinomial draws.

Applying this method to data from a systematic review of GBS carriage and disease, our analysis confirms longstanding clinical insight that GBS serotypes III and Ia pose the greatest risk of neonatal disease when carried by mothers, and that neonatal invasive infections caused by serotype III are the most likely to present as meningitis.^17–19^ Because this evidence preceded discovery of now-prevalent serotypes, the risks of serotype replacement under differing vaccine formulations have to date been uncertain.^7^ The enhanced association of serotypes III and Ia with invasive potential in the later neonatal period underscores the potential for GBS vaccination to reduce the burden of LOD, which cannot be prevented by screening-based intrapartum antimicrobial prophylaxis. We found serotypes VII and IX/NT-GBS to pose relatively limited risk of disease when carried by mothers. Although serotype VI has not historically been a prevalent cause of disease, its invasiveness appears on par with that of hexavalent vaccine serotypes other than III, suggesting serotype VI may be important to monitor in postlicensure surveillance. Taken together, our findings suggest non-vaccine types are unlikely to offset vaccine-preventable neonatal disease burden, even if they replace vaccine-targeted serotypes in carriage.

It is important to note that anticapsular vaccines may have impacts on GBS population structure beyond serotype replacement. The emergence of virulent vaccine-escape lineages by recombination constitutes an important threat foreshadowed by evolutionary responses of *S. pneumoniae* to PCVs.^32,33^ A majority of recombination events in GBS has involved genes encoding surface protein antigens and capsular biosynthesis proteins, suggesting immune escape via horizontal gene transfer.^34^ The acquisition of serotype IV capsule by clonal complex (CC) 17 lineages historically belonging to serotype III illustrates the capacity for invasive lineages to escape immunity via capsular switching. Expression of ≥7 distinct serotypes in CC1 isolates obtained from surveillance in Toronto underscores the frequency of serotype-switching events in GBS, and the need to understand the contribution of noncapsular antigens to epidemiologic parameters such as invasiveness.^35–37^ While our model can be extended to incorporate both clone and serotype as determinants of epidemiologic properties, molecular typing information in GBS is only beginning to become routinely available.^38^

The method we propose has value for analyses of multiple antigenically-diverse pathogens beyond GBS. Assessments of serotype replacement risk in pneumococcal disease remain important as vaccine manufacturers develop extended-valency anticapsular vaccines for *S. pneumoniae*. Increasing attention to the impact of PCVs on non-invasive pneumonia also underscores the need for determinations of the role of serotype in such disease.^39^ The method is also applicable to analyses of differential pathogenicity of *emm* types in group A Streptococcus, for which vaccines are currently in development;^40,41^ knowledge of *emm* type associations with carriage and disease states remains limited.^42,43^ Similarly, our approach may provide a strategy for understanding epithelial tropisms of the many human papillomavirus genotypes, and their role in cancer progression.^44^

Several features of our method can be extended. While not available for this analysis, prior distributions of both serotype frequencies and measures of pathogenicity can be incorporated into the calculation of posterior probabilities. For instance, phenotypic aspects of *S. pneumoniae* such as capsule width, molecular content, and charge predict serotype associations with carriage prevalence as well as risk of invasive infection,^45^ death,^25^ and otitis media;^15,46^ estimates of such associations could provide prior distributions on prevalence or invasiveness parameters for rare serotypes. Whereas we have assumed statistically-independent distributions of serotypes in carriage and disease within each setting, the model could also be extended to account for Dirichlet prior distributions of serotypes by geographic setting in addition to carriage or disease status.

In conclusion, we have provided a flexible empirical Bayes method for the estimation of serotype associations with carriage and disease states in meta-analyses. This method removes the need for ad-hoc continuity corrections that are commonly undertaken in such analyses due to sparse observations. Simulation studies reveal the method performs well under realistic conditions, providing 80% or greater coverage probability for multinomial probabilities from which we estimate measures of association. Applying this analytic framework, we identify low pathogenicity of non-vaccine GBS serotypes relative to serotype III, suggesting limited risk for vaccine-driven serotype replacement to offset the burden of vaccine-preventable GBS infections in neonates.

## SEARCH STRATEGY

We searched the following string in titles and abstracts of articles indexed in PubMed between 1 January, 2016 and 22 April, 2018 to identify relevant studies not included in previous reviews:

((“group b streptococc*”) OR (“streptococcus agalactiae”)) AND ((coloni* OR carria* OR carrie* OR carry*) AND (diseas* OR infect* OR invas* OR mening* OR sepsis OR septic*)) AND (serotyp* OR capsul* OR cps*)

We included studies that:

1. Were written in English, Spanish, German, or French.
2. Conducted serotyping by molecular or phenotypic assays for, at minimum, serotypes Ia, Ib, II, III, IV, and V, and reported the total number of isolates tested; this led to the exclusion of studies before 1999. We recorded the total number of isolates tested and the numbers identified as serotypes Ia, Ib, II, III, IV, V, VI, VII, VIII, and IX or NT-GBS; we pooled serotype IX and NT-GBS to accommodate data from studies undertaken before identification of serotype IX.
3. Described the population, setting, and study design: We excluded studies selecting patients based on microbiological findings post-hoc (e.g. studies limited to disease caused by a predefined sets clones or serotypes), and studies within which it was not possible to associate isolates with the specific colonization or disease outcomes under consideration (e.g. studies that did not specify whether patients were symptomatic or asymptomatic).

For analyses of neonatal invasive potential of serotypes, we included studies presenting “paired” sets of carriage and disease isolates, defined as isolates obtained from epidemiologically-linked individuals (i.e., mother-newborn dyads), or from geographically- and temporally-matched samples of neonatal disease cases and asymptomatically-colonized pregnant women. For analyses of serotype associations with disease manifestation and clinical outcome, we included studies presenting frequencies of individual serotypes in samples of all neonatal sepsis and meningitis cases, and in samples of all fatal and non-fatal neonatal invasive disease cases.

When studies presented data from multiple settings or populations, we abstracted data at the finest demographic and geographic resolution permitted by available metadata. We excluded articles presenting duplicate data from the same original study.

## SUPPLEMENTAL REFERENCES

The references listed here comprise all articles included in the review, matching the reference numbers listed in **Table 1** and **Table 2**.

## Acknowledgements

The authors thank Dr. Ruth Lynfield for helpful comments on an earlier version of the manuscript.

